# Rising through the Ranks: Seasonal and Diel Patterns of Marine Viruses

**DOI:** 10.1101/2020.05.07.082883

**Authors:** Gur Hevroni, José Flores-Uribe, Oded Béjà, Alon Philosof

## Abstract

Virus-microbe interactions have been studied in great molecular details for many years in cultured model systems, yielding a plethora of knowledge on how viruses use and manipulate host machinery. Since the advent of molecular techniques and high-throughput sequencing, viruses have been deemed the most abundant organisms on earth and methods such as co-occurrence, nucleotide composition and other statistical frameworks have been widely used to infer virus-microbe interactions, overcoming the limitations of culturing methods. However, their accuracy and relevance is still debatable, as co-occurrence does not necessarily mean interaction. Here, we introduce an ecological perspective of marine viral communities and potential interaction with their hosts, using analyses that make no prior assumptions on specific virus-host pairs. By size fractioning water samples into “free viruses” and “microbes” (i.e. also viruses inside or attached to their hosts) and looking at how viral groups abundance changes over time along both fractions, we show that the viral community is undergoing a change in rank abundance across seasons, suggesting a seasonal succession of viruses in the Red Sea. We use abundance patterns in the different size fractions to classify viral populations, indicating potential diverse interactions with their hosts and potential differences in life history traits between major viral groups. Finally, we show hourly resolved variations of intracellular abundance of similar viral groups, which might indicate differences in their infection cycles or metabolic capacities.

## Introduction

Viruses of marine microorganisms outnumber their hosts and are considered the most abundant biological entities in the ocean (1). They comprise the largest reservoir of genetic diversity in the oceans, and they are major participants in oceanic biotic and abiotic processes (2–7). It is estimated that ∼50% of marine microbial production is mediated by virus-induced release of dissolved organic matter or ‘viral shunt’ (8, 9). Viruses are suggested to alter microbial primary and secondary production (10–12), impact the population dynamics and diversity of microbial communities (13), and play an indispensable role in marine biogeochemical fluxes (14). Virus proliferation strongly depends on host metabolism (15), often by manipulating the host’s metabolic and transcriptional machinery using auxiliary metabolic genes (AMGs) (5, 16, 17). Additionally, recent environmental studies have reported several viruses infecting marine bacteria to be expressing AMGs in diurnal patterns, coupled with their host metabolism and reproduction cycle (18–22). Relying on their host for propagation, virus abundance is predicted to follow that of their hosts (“Kill-the-Winner” model (23), and the “Bank” model (24)). These models predict that in any given environment, a small fraction of viruses is highly abundant while the rest are rarer, waiting for the right conditions (i.e. host) to infect their hosts. Such abundance patterns are typical of microbial and viral communities (16, 24, 25), and their rank abundance distribution often fits a log-normal curve (26). While these concepts have been useful for describing the distribution of viruses in a given sample, our perception of the relationship between temporal abundance variation of marine viruses and host interaction remains largely obscure. Currently only a few environmental studies of diel patterns in marine viruses (18, 19, 22) and seasonality effects on viral communities (21, 27–32) have been reported, and none of these incorporate both diel and seasonal time-scales.

To resolve whether seasonal and diel patterns exist in the viral community (specifically, double-strand DNA viruses of marine prokaryotes) and to study the link between viral abundance and the potential interaction with their host, we collected two 24-hour time-series of coastal water samples during two seasons. We separated the cellular fraction (gDNA or metagenome), free virus fraction (vDNA or virome), and RNA fraction (RNA or metatranscriptome. Fig. S1b), allowing us to look at these fractions in both seasonal and diel time-scales. We used the Bank model (24) as a framework to classify viral active or inactive state based on their seasonal abundance across sample types (i.e. RNA, gDNA or vDNA), that is, classifying the abundant viruses as the “Active” group, and the rest as the “Bank” group (“inactive”) (Fig. 1a). By applying this framework, we show that most of the seasonally abundant viruses (Active) go through a seasonal change in abundance rank and that the high similarity in viral richness across seasons is largely derived from low abundance viruses (Bank). Furthermore, we assigned a taxonomic and ecological classification to hundreds of viral contigs with different patterns of abundance in the viral and cellular fractions. This classification indicates that similar seasonal abundance patterns of free viruses do not directly translate to similar abundance of these viruses inside their hosts (cellular samples), in contrast to what is often implied in viral metagenomic (i.e. viromes) studies. Finally, we observed many virus-host interactions (predicted by virus abundance in the cellular and RNA samples) having a differential signature between light and dark hours of the day. Such light-dependent interactions are prevalent in cyanophage populations (viruses infecting *Cyanobacteria*), but also include several viral populations predicted to infect a heterotrophic host.

**Fig. 1.**
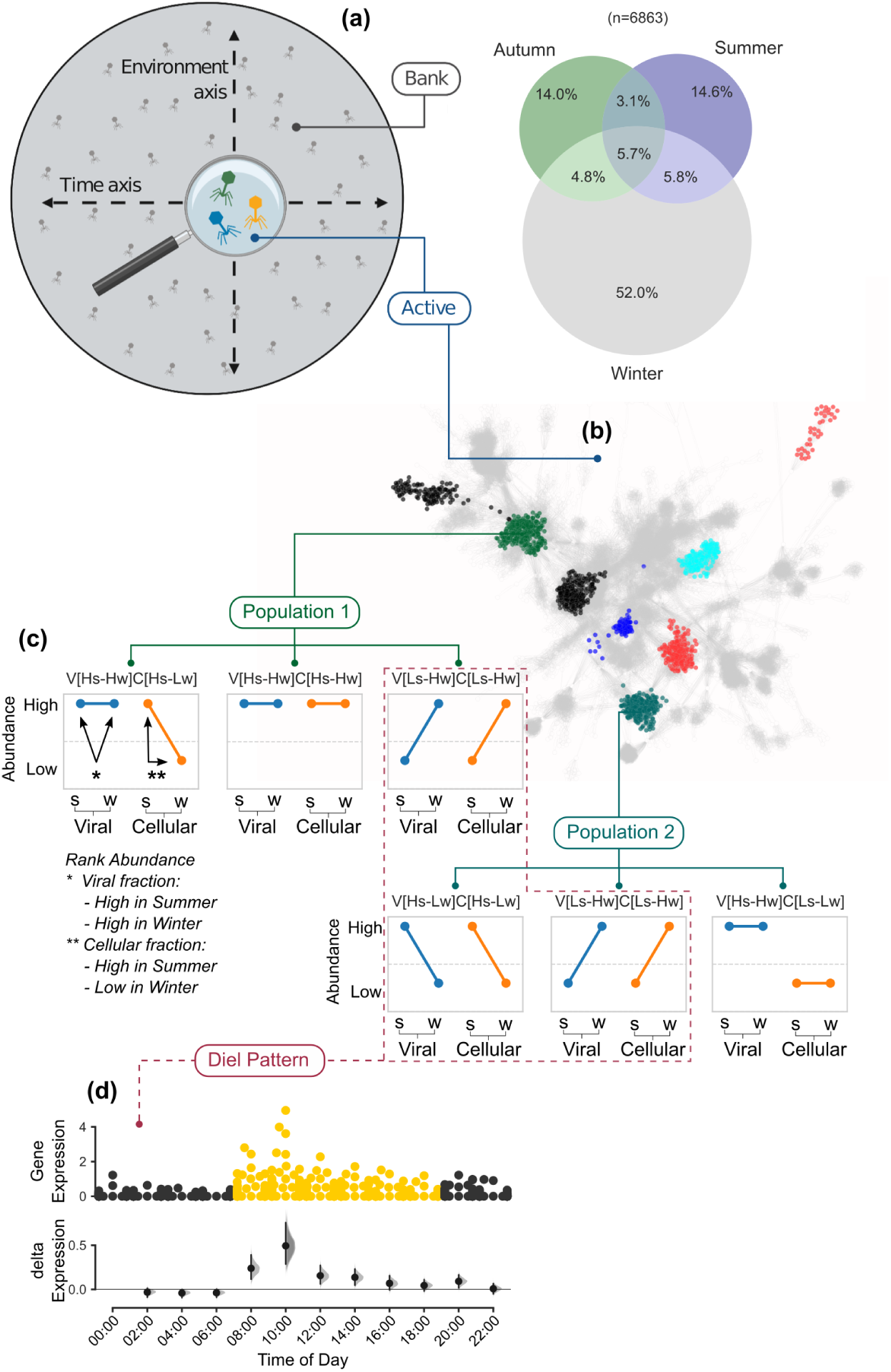
Classifying Viruses by seasonal abundance patterns. (a) Using the Bank Model (24) as a theoretical framework, the viral contigs were divided into an Active group and a Bank group. The venn diagram shows the percentage of shared contigs from the Active group in the viral fraction across seasons (b) Contigs from the Active group were annotated and clustered into populations using gene sharing networks (36). Each node is a viral genome/contig. Clusters color represent examples of distinct populations. (c) The rank abundance of each viral contig in the viral and cellular DNA samples was used to classify its seasonal abundance patterns (“rank-state”). Rank-states are annotated as follows: S (Summer); W (Winter); H (High ranks - Active group); L (Low ranks - Bank group); V (Virome samples) and C (Cellular metagenome samples). In each rank-state plot the left connected line (blue) represents the rank of a virus in the free-viral fraction in the summer (s) and winter (w) samples. Similarly, the connected line on the right (orange) represents the virus abundance in the intracellular fraction. As schematically shown for Populations 1 and 2 - a taxonomically cohesive population can have a diverse range of rank-states, while taxonomically distinct contigs can share the same rank-states. (d) Diel expression patterns of viral clusters can be linked to a rank-state. Each viral cluster was tested for intracellular light-dark differential signal (in the cellular or RNA fractions), showing that intracellular diel signal was mostly associated with viruses in strongly seasonal rank-states (V[Hs-Lw]C[Hs-Lw] and V[Ls-Hw]C[Ls-Lw]). X-axis, hours of the day when a sample was collected and the number of viral contigs expressed; upper panel, abundance distribution of viral contigs by time point (each point represents a contig); point color represent samples collected in the dark (black) and light (yellow); y-axis, contig abundance (RPKM); lower panel, estimation plot (37) displaying the effect size as a 95% confidence interval (1,000 bootstraps) of the mean differences between each time point compared against midnight as a reference group.

## Results and discussion

Two sets of 24-hour time-series samples with two hour intervals between samples were collected for this study. The first set was collected during the summertime (11-12 August, 2015) when the water column was stratified and the second was collected during late winter (7-8 February, 2016) when the upper water column was mixed. In addition, we used metagenomic data from a diel sample of Red Sea pelagic water collected during fall time (October, 2012) (33) (Fig. S1a). Assembled contigs (length ≥ 5kb) from all 48 DNA samples (cellular and viral fractions) were filtered for predicted viral contigs using two virus classifiers (see Extended Information). A total of 32,496 contigs were assigned as viral by both algorithms (minimum probability cutoff 0.75) and were mostly enriched in the viromes compared to the metagenomes (Fig. S2a). These contigs were considered of viral origin for downstream analysis. The rank abundance distribution of the viral contigs best fitted a log-normal distribution in all the datasets (34) (i.e. different sample types and seasons. (Fig. S2b) and Table S1), indicative of a relatively small number of viruses with high abundance, while the majority have a very low abundance (in agreement with the Bank model and other reports (18, 21, 24, 25)). We designated the most abundant viral contigs as the Active group and the rest as the Bank group (abundance threshold was set at 80% mapped reads from the total reads. See Extended Information). This classification revealed that the Active group is seasonally disparate both as free viruses and as viruses in the intracellular samples (Fig. 1a, venn diagram and Fig. S3, right column)). In contrast, when considering the viral contigs that are present across seasons (reads per kilobase per million mapped reads [RPKM] > 1), it appears that the seasonal communities share a large proportion of contigs in the viral and cellular samples but less so in the RNA fraction (77.2%, 45.0% and 11.9% in the viral, cellular and RNA samples, respectively. Fig. S3, left column). This suggests that the rank distribution’s long right-tail of low abundance viruses (Bank) is responsible for most of the viruses that are present across seasons (32), while the top-ranking members of the viral community are undergoing a seasonal succession (Fig. 1a). This seasonal change of ranks, together with the log-normal distribution and highly shared fraction of low abundance viruses, supports the Bank model where low abundance viruses in one environment (stratified summer water in our case) increase in abundance as the environment changes (e.g. mixed winter water) and their host abundance increases (18, 24). Furthermore, the considerable winter increase in the number of viruses contributing to the Active group, suggests an increase in both the diversity and abundance of potential hosts (14.6% unique contigs in the summer vs. 52% in the winter viral fraction Fig. S3). A similar mechanism, where high seasonal host diversity generates high viral diversity has been recently hypothesized to be in effect in the Arctic marine environment (35). Our observations suggest that seasonal variations in host diversity could drive an increase in viral diversity in a coastal and more temperate marine environment, and are not confined to the Arctic.

We classified the Active viral group into viral clusters (VC) based on gene-sharing network (36) (Fig. 1b and Extended Information) and a taxonomic annotation was assigned based on the identity of Refseq viral genomes found in that cluster, or according to the closest Refseq genomes that represented a distinct hub. We kept the annotation at the viral genus level (38) (e.g. cyanopodovirus, SAR11 virus, etc.) and considered highly connected VCs with the same taxonomic annotation as “viral populations”. Most of the VCs did not cluster with a Refseq representative and were classified as “Uncultured virus” in our data (Table S2), accentuating the high proportion of yet uncharacterized viruses in the environment (2, 18, 25, 35). A closer look at the abundance of different viral populations across seasons and sample types reveals different patterns for distinct populations. For example, cyanophages, one of the most abundant viral groups in our data, was grouped into four populations according to their viral families, *Cyanomyoviridae, Cyanopodoviridae*, and two *Cyanosiphoviridae* populations (Fig.2). The cyanomyovirus population (300 contigs) was abundant in both viral and cellular samples in all seasons. The cyanopodovirus population (220 contigs) was abundant in the viral samples across seasons, while in the cellular samples it was highly abundant in the summer, but mostly absent from the autumn and winter samples. One cyanosiphovirus population (Fig.2, lower right, 175 contigs) was abundant in the viral samples across all seasons but almost absent from the cellular samples. In contrast, another cyanosiphovirus population (Fig.2, upper right, 34 contigs) showed a distinct sub-population difference in abundance between summer and winter, where VCs 251_0 and 550_0 were abundant in the summer samples and VCs 281_0 and 419_0 in the winter samples (in both viral and cellular samples). The observed variability in the seasonal abundance of these closely related viral populations highlights the different life history traits of these populations and sub-populations in terms of possible host range and life style, suggesting potential adaptations to changing host landscapes. Indeed, a comparison of the most abundant microbial taxonomic groups show considerable differences between summer and winter (Fig. S4).

**Fig. 2.**
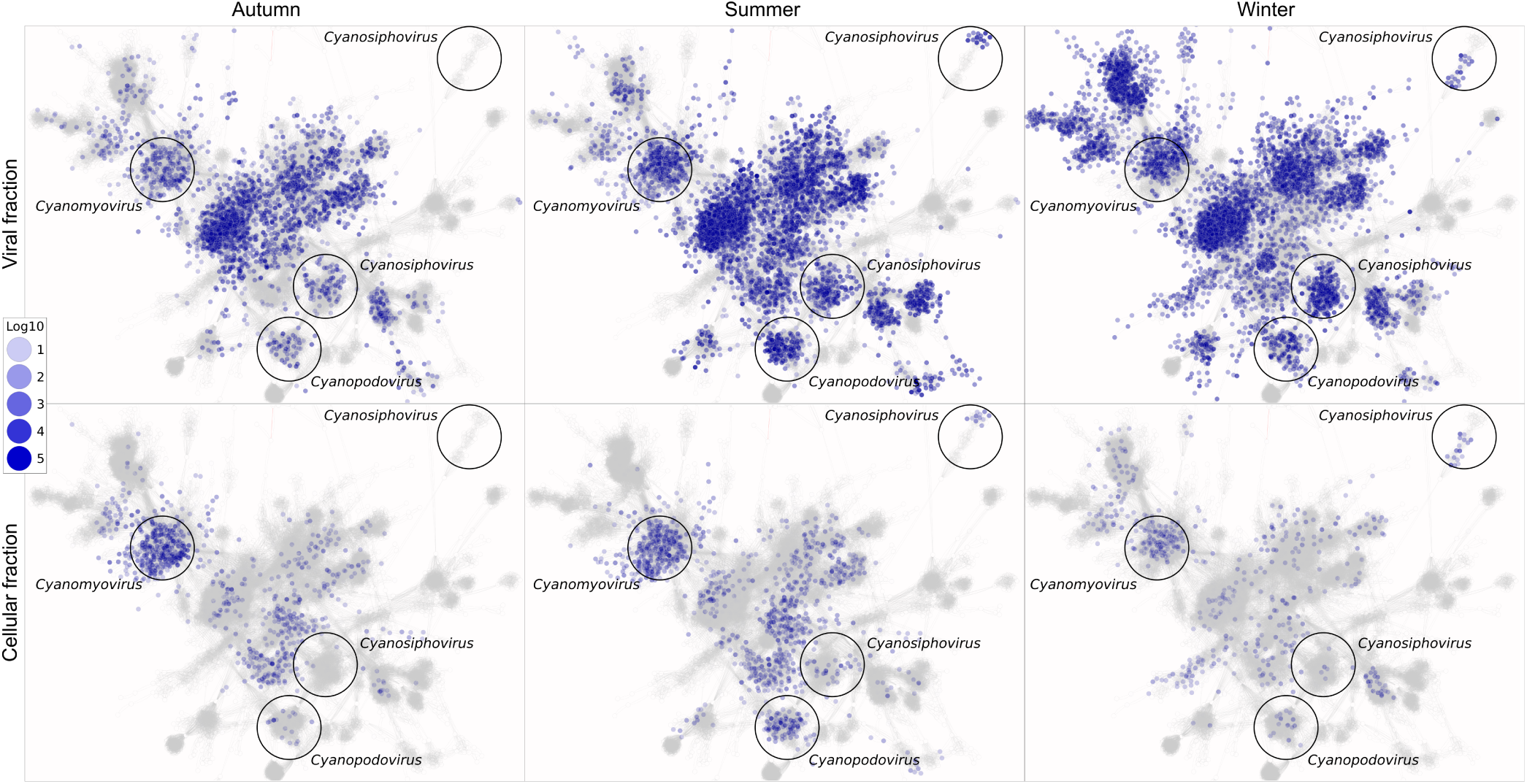
Seasonal virus abundance overlaid on a gene sharing network. Each node is a viral genome/contig. Color intensity represents log abundance. Top: Viral fraction. Bottom: Cellular fraction. Labeled are cyanophage groups discussed in the text.

To systematically assign an ecologically relevant classification to viruses based on seasonal abundance we used the position of each viral contig along a scaled rank abundance (between 0 and 1) in each of the different viral and cellular DNA samples (Fig. 1c). Based on the concepts of the Bank model (24), we looked for seasonal transition of each viral contig between the Active and the Bank groups (i.e. a move from the top 20% most abundant contigs in one season to a lower abundance in another season and vice versa). We term these abundance-sample combinations “rank-states” and annotated them as follows: S (Summer); W (Winter); H (High ranks Active group); L (Low ranks Bank group); V (Virome samples) and C (Cellular metagenome samples). For example, high abundance of a group in the viral sample in both summer and winter is annotated as **V**iral[**H**igh**s**ummer-**H**igh**w**inter] and abbreviated as V[Hs-Hw]. Furthermore, to differentiate between the Active and Bank groups, we defined a seasonal change in rank group (i.e. from Active to Bank and vice versa) as a minimum of 20% change on the rank scale and one order of magnitude change in abundance. The combination of two seasons (summer, winter) and two sample types (viral fraction, cellular fraction) yields 16 possible combinations, representing the relationship between a virus abundance in the free viral fraction and its abundance when interacting with its host in the cellular fraction, in a changing environment (i.e. seasonal variation). Furthermore, the extent of rank change can also be deduced and, for example, contigs that undergo no change in rank across seasons are easily identified (Fig. 3). Classifying the Red Sea viral community using this framework reinforces the observation of various abundance patterns for different viral populations, while assigning each viral contig an ecological context (Table S3). In the summer samples (V[Hs-Lw]), 21.75% of the viral contigs were more abundant in the viral fraction in comparison to 43% in the winter samples (V[Ls-Hw]). In contrast, the abundance of viral contigs in the cellular fraction (i.e. C[Hs-Lw] and C[Ls-Hw]) represented 21.85% of the summer sample but only 5.53% of the winter samples (Fig. 3 and Table S4). These results further suggest that the abundance of free viruses alone (as measured in the viral fraction) does not directly correlate to their abundance in the cellular fraction and vice versa, and thus cannot reliably serve as a predictor for infection levels. Additionally, this approach can highlight interesting VCs with a V[Ls-Lw]C[Hs-Hw] rank-state (Fig. 3, bottom left) that show a remarkable pattern of medium abundance in the viral samples, with considerable seasonal change in rank, in contrast to a high stable abundance in the cellular fraction. As the majority of contigs in this rank-state appear to be of uncharacterized SAR11 viruses, the traits that enable this pattern are still unknown.

**Fig. 3.**
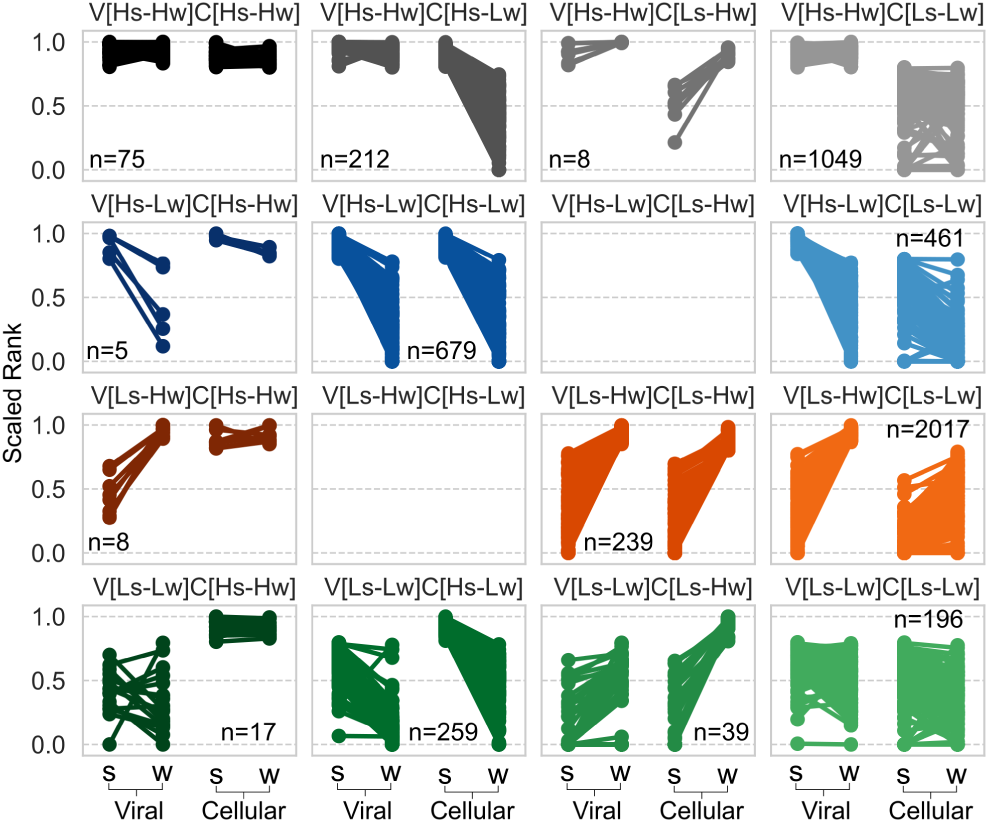
Representation of viral rank-states of contigs from the Active group. Including 16 possible rank-state combinations, from viruses that are highly abundant in both seasons and samples (V[Hs-Hw]C[Hs-Hw] n=75, top left), to those with lower abundance across seasons and samples (V[Ls-Lw]C[Ls-Lw] n=196, bottom right)

Diel cycles lie at the basis of ocean productivity as sunlight is the most readily available source of energy in the photic zone (39), driving microbial community-wide gene expression patterns (40). Many of the diel gene expression patterns of the microbial community in our samples (collected concurrently with the viral samples) presented a strong correlation with a daylight cycle and weak seasonal hierarchical clustering. This correlation was robust across seasons and functions of the microbial community, indicative of potential community-level trait-diel relationship and functional redundancy among these seasonally varying communities (41, 42) (Fig. **??**). Since virus proliferation strongly depends on its host metabolism (15), we followed the patterns in the microbial community to examine seasonality and daylight effects on the viral community intracellular presence (18). Generally, in the RNA fraction we observed an increase in the transcript abundance of the cyanopodovirus population at 08:00 (first sample collected in light) and a peak at 10:00 in the summer sample with a slight shift in the winter samples (Fig. 4c, top). This peak included the expression of phage structural genes and key cyanophage AMGs such as *psbA* (subunit of Photosystem II) and *hli* (a putative stabilizing protein for photosystem II), indicating active infection. Abundance of cyanopodoviruses in the cellular fraction peaked at 10:00 in the summer and was again shifted in the winter sample (Fig. 4d, top), consistent with the earlier peak of transcription of viral genes observed in the RNA fraction. In addition, a clear difference in the diel pattern can be observed between rank-states (seasons), despite the fact that they originate from the same viral population (Fig. 4, top). The cyanomyovirus (Fig. S7) population’s transcript abundance also increased with the onset of light and fluctuated during the day, with most genes at their highest around 18:00. Expressed viral genes included structural genes as well as AMGs such as *psbA, hli, CP12* (Calvin cycle inhibitor) and *phoH* (phosphate inducible protein). The abundance patterns of cyanomyoviruses in the cellular fraction displayed multiple peaks, most pronounced at 10:00-12:00 and at 16:00-18:00 (Fig. S7). The cyanosiphovirus population’s transcript abundance peaked at midnight and noon, while their cellular abundance increased during daytime, peaking at noon (Fig. S7). To understand the finer details of these patterns and their seasonality, we examined the light-dark intracellular abundance patterns of viral clusters with a minimum of 10 open reading frames (time points classified as light or dark samples. Total of 441 VCs with *ge* 1 RPKM in the light or dark samples). This analysis revealed 97 VCs (22%) with at least one rank-state that displayed differential light-dark abundance (Mann-Whitney U test, p < 0.05. Fig. S6a and Table S5). Most of the light-dark differential signal in either the cellular or RNA fractions came from 69.57% of these VCs, specifically from contigs in only four rank-states (V[Ls-Hw]C[Ls-Lw], V[Hs-Lw]C[Hs-Lw], V[Ls-Lw]C[Hs-Lw], V[Ls-Hw]C[Ls-Hw]). Moreover, most of the RNA signal came from only two rank-states of viruses with distinct summer (V[Hs-Lw]C[Hs-Lw]) and winter (V[Ls-Hw]C[Ls-Hw]) abundance patterns. This signal was very low or undetectable in other rank-states, indicating that most of the detected viral RNA transcripts are from viruses that show a strong seasonal pattern, both extra- and intracellularly (Table S5). When examining the seasonal patterns of the microbial groups in the cellular and RNA fractions, it appears that the seasonal microbial communities are largely different from one another, sharing only 19.5% of the contigs (23,219 shared contigs). Additionally, while the *Synechococcus* population is highly abundant and active across our seasonal samples, changes in the dominance of unicellular picoeukarya algae in the winter and *Prochlorococcus* in the summer (Fig. S4), is consistent with previous reports in this region (43, 44). Indeed, as would be expected by these patterns of potential microbial hosts, a large proportion of the VCs with significant light-dark differential abundance was part of the cyanophages group (488 contigs in 16 VCs and 9 rank-states. Fig. S6b and Table S5). Some cyanophages have been shown to have a light dependent lytic infection cycle, where the infection is initiated in the early morning and progresses throughout the day when energy fluxes are at their peak (through photosynthetic activity), promoting viral propagation (20, 45–51). Recent work has also shown that some Prochlorococcus viruses are capable of adsorption and even replication in the dark (51). The cellular and RNA diel abundance of some of the cyanophage populations in our data show a pattern that peaks during light hours, suggesting that the infection of these cyanophages indeed follows diel energy fluxes as previously suggested (18, 20, 21, 50, 51). Summer-abundant cyanophages (in rank-state V[Hs-Lw]C[Hs-Lw]) show an increase in intra-cellular abundance during light hours in all three families. In winter-abundant cyanophages (V[Ls-Hw]C[Ls-Hw]) similar daytime increase was observed for cyanopodoviruses and cyanomyoviruses, but not for cyanosiphoviruses (Fig. 4 cyanopodoviruses, and Fig. S7 cyanomyoviruses and cyanosiphoviruses). The variation in intracellular abundance patterns by viruses infecting similar hosts might indicate differences in the infection cycle or the metabolic capabilities of these viruses as has been recently described for three different cultured *Prochlorococcus* viruses (51, 52). That is, based on our results we hypothesize that a high diversity exists in the timing and rate of various key aspects of infection such as absorption, virion production, and lysis for these viruses in their natural environment.

**Fig. 4.**
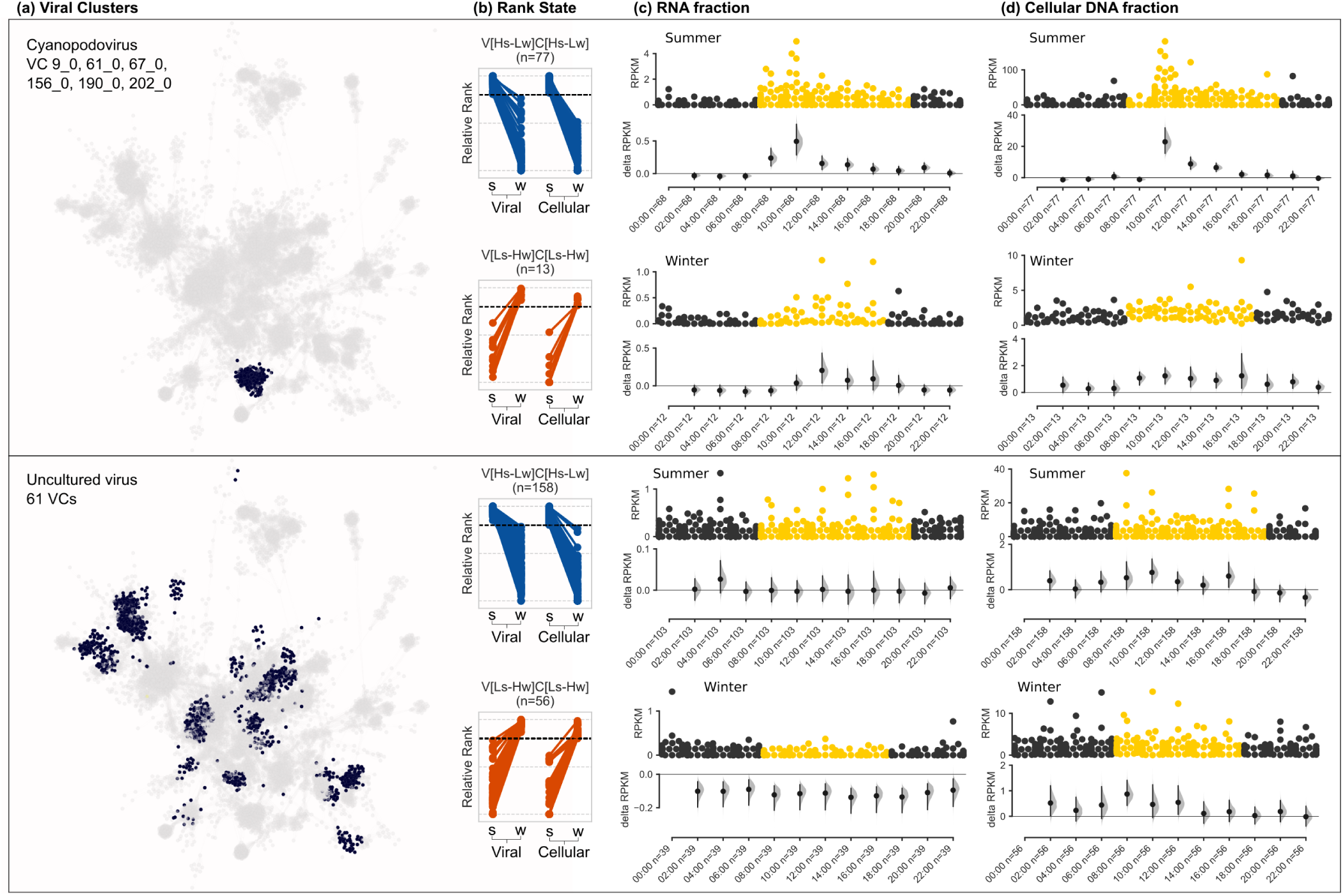
Intracellular diel abundance patterns of cyanopodoviruses (top) and uncultured virus populations (bottom). (a) Viral clusters of contigs from three cyanophage families grouped using gene-sharing network (Bin Jang et al. 2019). (b) Seasonal rank-state abundance patterns of selected contigs from the VC on the left that displayed a differential light-dark signal (Mann-Whitney U test, p < 0.05). (c-d) Diel distribution of intracellular RNA (c) and DNA (c) of viral contigs from a specific VC and rank-state (left); x-axis, hours of the day when a sample was collected and the number of viral contigs expressed; upper panel, abundance distribution of viral contigs by time point (each point represents a contig); point color represent samples collected in the dark (black) and light (yellow); y-axis, contig abundance (RPKM); lower panel, estimation plot (37) displaying the effect size as a 95% confidence interval (1,000 bootstraps) of the mean differences between each time point compared against midnight as a reference group.

The contrast between the detection of viral transcripts in the different cyanophage families, where the podo- and myoviruses show a clear transcriptional increase at the onset of light, as opposed to the less clear pattern in the siphoviruses, could result from lack of AMGs in the latter (based on currently described genomes). Further manual inspection of several complete cyanosiphovirus genomes in our samples (length ≥ 40kb) confirmed the absence of known AMGs in these genomes. However, this hypothesis cannot be validated using our data and requires additional experimental validation. These results signify that while most cyanophage populations display a clear intracellular increase in abundance during daytime, the hourly resolved variations might indicate differences in their infection cycles or capacities to tinker host metabolism with AMGs.

Interestingly, our approach also detected a large group of uncultured viruses (947 contigs in 61 VCs and 9 rank-states) with light-dark differential abundance signal (Fig. S6b and Table S5). Many of these viruses putatively infect heterotrophic microbial hosts (based on proximity to a Ref-Seq viral genome in the sequence similarity network. Fig. S8). This large group of viruses exhibits diverse abundance patterns and potentially different diel-dependent life history traits. For example, contigs in this group with a high summer presence in both the viral and cellular fractions (V[Hs-Lw]C[Hs-Lw]) showed increase in intracellular abundance during light hours (a handful in the RNA and the majority in the DNA fraction), while with their winter counterparts (V[Ls-Hw]C[Ls-Hw]) such an increase was only observed in the cellular DNA and not in the RNA (Fig. 4, bottom). This result extends the previously described diel and seasonal activity repertoire of viruses of heterotrophic microbial hosts in the marine environment (18, 19).

Differential light-dark virus-host interaction is thus prevalent among viruses in the photic marine environment and is not restricted to cyanophages. However, while the uncultured virus group is larger than the cyanophages group, the prevalence of contigs with detectable differential light-dark RNA signal was more pronounced in the latter (Table S5). In addition, we observed low abundance of transcripts from viruses of heterotrophic bacteria (SAR116 as a prime example, Fig. S7, bottom) despite the strong transcriptional diel patterns of many of their potential heterotrophic bacteria hosts in this study (Fig. S5 and Table S6). Our results are consistent with previous reports which mostly detected cyanophage transcripts in environmental metatranscriptomic samples (18), indicating their high levels of intracellular transcription in comparison to highly abundant heterotroph viruses (e.g. SAR11 and SAR116 viruses). In addition, viruses infecting heterotrophic microbes such as SAR11, SAR116, MGII Euryarchaea (33, 53, 54) have been previously reported to have lower intracellular signal (33, 55). Notwithstanding the high abundance of free viruses, these results indicate that the fraction of actively infected heterotrophic hosts is low in our samples, especially in comparison to cyanobacteria and their viruses. It is possible that the low signal in the RNA and cellular fractions is due to an experimental bias, or to the fact that they operate on longer and slower life cycles (53, 54). However, our observations were consistent across multiple seasons, samples and technical replicates. Furthermore, our ability to detect transcription of the hosts of these viral groups and to detect high levels of abundance in the viral fraction indicates no bias against them in the computational analyses. The discrepancy between the infection levels of cyanophages and that of viruses of heterotrophs suggests a fundamental difference in lifestyle and infection dynamics between these groups. In light of this, it is most intriguing that viruses of heterotrophs remain in high proportions in the Bank, and future research is needed for understanding their underlying infection dynamics in the marine environment.

## Conclusions

Here we elucidated previously unknown complex seasonal and diel patterns of abundance and activity of marine viruses of the photic zone. We used the Bank model (24) as a conceptual framework where the most abundant viruses are considered Active and the remaining majority as Bank. In addition, we classified viruses based on their seasonal changes in ranks. The Bank group persisted across seasons, while most of the Active group showed seasonal change in rank abundance in either the viral or cellular fractions. We speculate that the seasonal viral response is facilitated by a stable Bank population that is readily activated upon increase in abundance of a potential host, in accordance with Kill-the-Winner and Bank models (23, 24). A negative correlation between virus production (burst size) and virus survival have been previously demonstrated even between viruses of *E. Coli* (56). Thus, beyond the straightforward implications regarding the infection cycle of these viral groups (Bank and Active) in the marine environment, questions arise regarding their life history traits and challenge the prevailing view of virus decay rates (9, 57). Specifically, the mechanisms that allow for viral groups to persist in the Bank population despite apparent low infection levels are currently not being considered in most models. The detection of multiple rank-states in taxonomically similar viral populations suggests that the overall population-level response of viruses is resilient to changes in host composition (58). This supports the theoretical observation that virus-host interactions cannot be easily deducted from linear relationship in their abundance in surface water samples (1). Our observed diversity in diel patterns of viral transcripts and intracellular abundance, within both VCs and rank-states, provide environmental support to recent culture based evidence (51, 52) and suggests that it is more widespread. Our observations in the Red Sea of seasonal changes in the ranks of the most abundant virus groups are supported by a previous long-term study in a freshwater lake (31), yet contrasts a recent report of mostly stable viral populations in a five-year time-series study at the San Pedro Ocean Time Series (SPOT) (32). These contrasting observations could potentially result from different hydrodynamic properties governing each site. The Red Sea is mostly restricted to input from other marine sources (its only source is through the shallow Straits of Tiran), and the plankton community composition is governed by seasonal cycles of up-welling and deep convective mixing that bring nutrients to the otherwise oligotrophic surface water (59). Such properties could be more analogous to those of the lake than the SPOT site. Additionally, our data suggests that high viral abundance in the viral fraction might not necessarily translate into high infection levels, as is evident in the case of many heterotrophic viruses with almost no detectable intracellular presence, despite being highly abundant in the viral fraction. This could suggest a consistently low production of virions (below detection levels), allochthonous inputs, extremely slow decay rates or another as yet unknown mechanism supporting this intra- and extracellular abundance discrepancy.

The approach used in this study facilitated the discovery of several co-existing viral populations that exhibit various seasonal and diel patterns. The prevalence of differential diel light-dark abundance patterns of viral contigs across seasons might indicate the convergence of infection and propagation strategies employed by viruses as a function of the recurrent diel metabolic patterns of their hosts. Furthermore, the repertoire of possible viral lifestyles and infection dynamics, manifested in rank-states in this study, suggest underlying differences within major virus groups in the marine environment. Additional fundamental differences between viruses of marine photosynthetic microbes and viruses of heterotrophs appear to exist and further investigation is needed to understand how life history traits differ between these groups and whether they can explain the patterns we observed. The VCs and rank-states reported here can serve as potential targets for such future research, which could be fundamental for understanding the contribution of viruses to the structure of microbial assemblages and the resulting impact on the dynamics of viral shunt, nutrient cycles and the marine food web.

## Data availability

The assembled viral contigs, along with raw sequence data, are available from the European Nucleotide Archive (ENA) under accession PRJEB35627. Supplementary Tables can be downloaded from https://doi.org/10.17605/osf.io/b74mt.

## Code availability

Custom code is available at: https://doi.org/10.17605/osf.io/b74mt

## ACKNOWLEDGEMENTS

The authors would like to thank The Interuniversity Institute for Marine Sciences in Eilat, Israel (IUI) for providing access to their pier as well as work space and the Genomics Centers at the Biomedical Core Facility and Lorry I. Lokey Interdisciplinary Center for Life Sciences and Engineering at the Technion for library preparation and sequencing. The authors would also like to thank Dan Fishman for samples preparation, Itai Sharon for help with metagenome assemblies, Sheila Roitman and Faris Salama for their help with sampling, and David Cohen of the Physics Department at the Technion for his help with the HPC ATLAS cluster used for the analyses. This work was funded by a European Commission ERC Advanced Grant (no. 321647), the Technion’s Lorry I. Lokey Interdisciplinary Center for Life Sciences and Engineering and the Russell Berrie Nanotechnology Institute, and the Louis and Lyra Richmond Memorial Chair in Life Sciences (to O.B.).

## Extended Information

### Sampling site

Evident seasonal succession patterns of microbial communities have been previously described in the Gulf of Aqaba (44, 59, 60), largely driven by its oligotrophic nature and annual stratification-mixing cycles (59, 61, 62). Additionally, changes in viral communities have also been observed between different seasons and and between stratified and mixed water column (30, 63– 66).

### Sample collection

Water samples were collected every two hours during a span of 24 hours, from the Interuniversity Institute (IUI) pier in Eilat from a depth of 2-3 meters (surface water). At each time point, two 20L containers of seawater were initially filtered through a GF/D glass microfiber filter (Whatman) with a nominal pore size of 2.7m to remove large eukaryotic cells. One 20L container was used to collect two fractions: cellular genomic DNA (cellular fraction or metagenome) and, after filtration, viral DNA (viral fraction, virome or free-viruses). The second container was used to collect RNA (RNA fraction or metatranscriptome). A similar set of samples was collected on August 11-12, 2015 (summer samples) and on February 7-8, 2016 (winter samples). The cellular DNA fraction was collected onto a 142 mm 0.22m Durapore filter (Millipore) using a peristaltic pump. DNA was extracted using the alkaline-lysis protocol. For the viral DNA, the flow through of the 0.22m filters was treated with iron-chloride (FeCl3) for the precipitation of viral particles, as described in (Poulos, John, and Sullivan 2018). The precipitate was collected on a 0.22m Durapore filter (Millipore) and washed with 10ml of Calcium Oxalate solution. The washed precipitate was further concentrated using a 100kDa Centricon filter (Millipore) and purified on a CsCl gradient followed by DNase treatment as described in (67). DNA extraction was performed using Wizard mini-columns (Promega) as described in (68). The meta-transcriptome fraction was collected onto a 142 mm 0.22m Durapore filter (Millipore) using a peristaltic pump. After collection, the filters were transferred immediately to a screw cap containing 1 ml of RNAlater (Ambion) and frozen in liquid nitrogen. Total handling time was less than 15 min. Total RNA extraction was done using the mirVana RNA isolation kit (Ambion), followed by DNA removal with Turbo DNase (Ambion) and cleanup using the RNeasy MinElute Kit (QIAGEN).

### Sample processing and sequencing

The DNA samples were sheared using Covaris E220 with the following parameters: 10% duty factor, 45s duration time, 200 cycles per burst, 175W peak incident power and a temperature of 6. The RNA samples were fragmented using a library preparation kit (NEBNext) with 5 minutes in 75. The mean fragment lengths (without adapters) of the DNA and RNA samples was 404 and 350 bp, respectively. None of the samples were amplified prior to sequencing-library preparation, nor were rRNA depletion protocols applied in order to avoid possible bias resulting from these steps (69) and loss of considerable amount of RNA during the rRNA depletion process. Libraries were constructed with 10ng of DNA per sample using NEBNext Ultra II DNA Library Prep Kit with 12 PCR cycles for the DNA samples, and 100 ng with NEBNext Ultra RNA Library Prep Kit with 15 PCR cycles for the RNA samples. All samples were paired-end (PE) sequenced at the Technion Sequencing center on Illumina Hiseq 2500, where the DNA samples sequenced with 2×125 bp, and the RNA samples with 2×100 bp.

### Quality control and assembly of short reads

Reads were trimmed using Trim Galore version 0.4.4 (https://github.com/FelixKrueger/TrimGalore) with default parameters. Data for the different 48 DNA samples (24 vDNA and 24 gDNA) were assembled separately using IDBA-UD (70) with default parameters.

### Annotation

The assembled contigs from all the different samples were concatenated in a single FASTA file and dereplicated to keep those 5000 bp using vsearch version 2.6.2 (71) with the options ‘–derep_fulllength –minseqlength 5000 –maxseqlength 5000000’. The de-replicated contigs were then screened using Mash version 2.0 (72) for similarities to PhiX (GenBank accession: HM775309.1), a common control used during Illumina sequencing runs which can result in spurious assemblies (73). Using the Mash results those contigs with similarities to PhiX were discarded from further analysis. Open reading frames (ORFs) of the remaining contigs were predicted by Prodigal version 2.6.3 (74) using the parameter ‘-p meta’. ORFs longer than 300 nt were dereplicated using vsearch with the options ‘–derep_fulllength –minse-qlength 300 –maxseqlength 5000000’. ORFs were taxonomically annotated using either BLASTn best-hit against NCBI nr, Diamond version 0.8.38 (75) ‘blastp’ best-hit to the protein sequences from the Prokaryotic Virus Orthologous Groups (pVOGs) database (76). ORFs appearing in the RNA data were assigned a functional category using eggNOG mapper version 1 (77, 78), with mapping mode “Diamond” and default parameters.

### Relative abundance estimations

The relative abundance of ORFs was calculated using Salmon version 0.8.2 (79). A total of 16,047,552 ORFs predicted from 3,878,543 assembled contigs and 3,224 Refseq genomes (bacteria, archaea, plastid, protozoa and viruses; accessed: 2017-02-02. Table S5) were dereplicated using vsearch ‘–derep_fulllength’ (10,893,032 unique ORFs) and used to create a Salmon index. The Refseq genomes used were selected by classifying the unassembled reads with Kraken version 0.10.4-beta (80) and a custom database composed by all Refseq available genomes (bacterial, archaeal, and viral; accessed: 2017-02-02). The list of taxa assigned to the reads was retrieved and processed to keep only those genomes not considered as sample-processing contamination sources (81). The ORFs abundance in the 72 datasets (metagenomes, metatranscriptomes and viromes) was quantified with the index using Salmon in the quasi-mapping mode with the following parameters ‘–meta –incompatPrior 0.0 – seqBias –gcBias –numAuxModelSamples 2500000 –numBootstraps 100 –validateMappings’. Quantification results were processed by tximport (version 1.10.0) (82), followed by normalization with edgeR (v3.24.2) (83). Reads per kilobase per million were calculated from the normalization results by the edgeR function ‘edgeR::rpkm’.

### Clustering diel gene expression patterns

Predicted ORFs were quantified as mentioned above (Salmon and tximport), followed by normalization with Bioconductor R package DESeq2 (84) by applying a regularized log transformation (rlog) to the Salmon counts data (baseMean 2). The normalized counts were clustered using Gaussian mixture models coupled with Dirichlet process (DP-GP) (85) with default parameters. The ORFs clustered into 18 and 23 clusters in the summer and winter datasets respectively, based on the similarity of their diel expression patterns. Mapping, binning and prediction of viral contigs Short reads were mapped to the the assembled contigs using Bowtie2 version 2.3.4.1 (86) and Samtools 1.3.1 (87). The mapping files were used to generate a depth coverage matrix using Metabat2 script ‘jgi_summarize_bam_contig_depths’ and binned with Metabat2 version 2.12.1 (88) with parameters -m 1500 -s 10000. Prediction of viral contigs was performed using two independent classifiers: (i) VirFinder (89), R package version 1.1 with default parameters and modEPV_k8.rda model for predicting both prokaryotic and eukaryotic viruses. (ii) MARVEL (90) version 0.1 with default parameters.

### Rank abundance model fitting

Viral contigs with length ≥ 5kb and abundance RPKM ≥ 1 were used to evaluate the rank abundance distribution by season and sample type, with Vegan R package version 2.5-3 (34), using the ‘radfit’ function with default parameters. The models were evaluated based on their Akaike information criterion (AIC) values, where the model with the lowest AIC was determined as “best fit”.

### Classification of viral contigs to Active or Bank groups

The most abundant contigs to which a total of 80% of the reads mapped back, were designated as the Active group (n=7,047). Contigs in this group represented 12.5% from the total number of contigs in the summer samples and 21.6% of the contigs in the winter samples. The remaining contigs were designated as the Bank group. The viral contigs in the Active group were used downstream for taxonomic annotation.

### Taxonomic classification of prokaryote viruses and network visualization

Taxonomic assignment of viral contigs from the Active group was performed with vConTACT 2.0 (36) on the CyVerse cyberinfrastructure platform (91). Viral contigs were clustered into viral clusters (VC) based on gene-sharing network, where each VC was assigned a taxonomic annotation based on the identity of Ref-seq viral genomes found in the same VC, or according to the closest Refseq genomes that represented a distinct hub (node degree » average node degree. The vConTACT analysis was performed with the following parameters: Protein clusters generation method: MCL, protein-protein similarity method: Diamond, VC generation method: ClusterONE, and Reference database: NCBI Bacterial and Archaeal Viral RefSeq V85 with ICTV + NCBI taxonomy. Network visualization was performed using Cytoscape version 3.7.1.

## Extended Figures

**Fig. S1.**
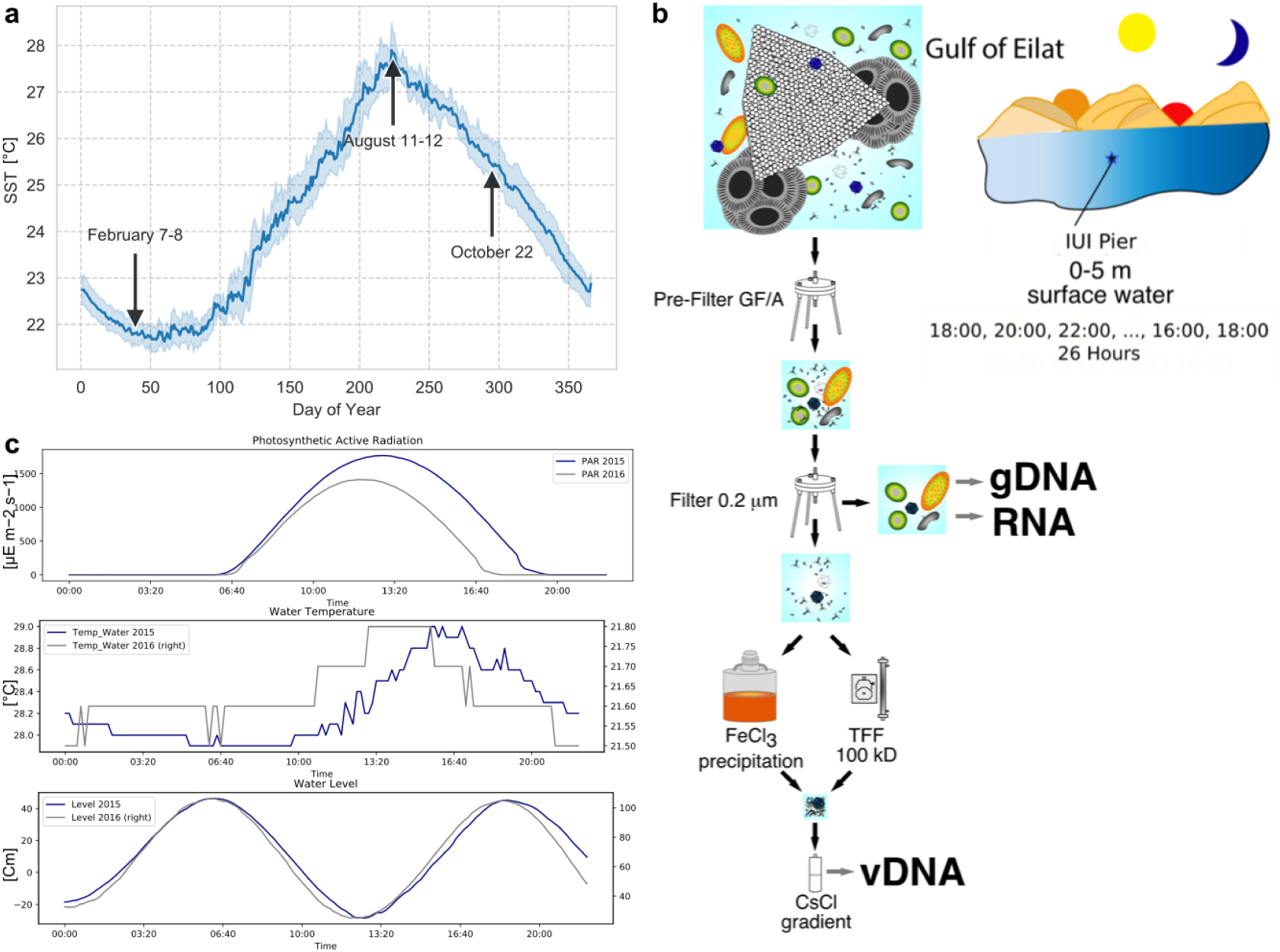
(a) Sea surface temperature (SST) measured at the IUI Pier from 2011-2019. x-axis: Day of Year, y-axis: mean SST with 95% confidence bands (1000 bootstraps). Arrows indicate dates of sampling mentioned in this study. (b) Sampling scheme. Sea water samples are every two hours, filtered and fractionated based on microorganisms size and nucleic acid content. (c) Physical parameters recorded at the IUI pier during days of sampling in August 2015 (blue) and February 2016 (grey). PAR, water temperature, and water level.

**Fig. S2.**
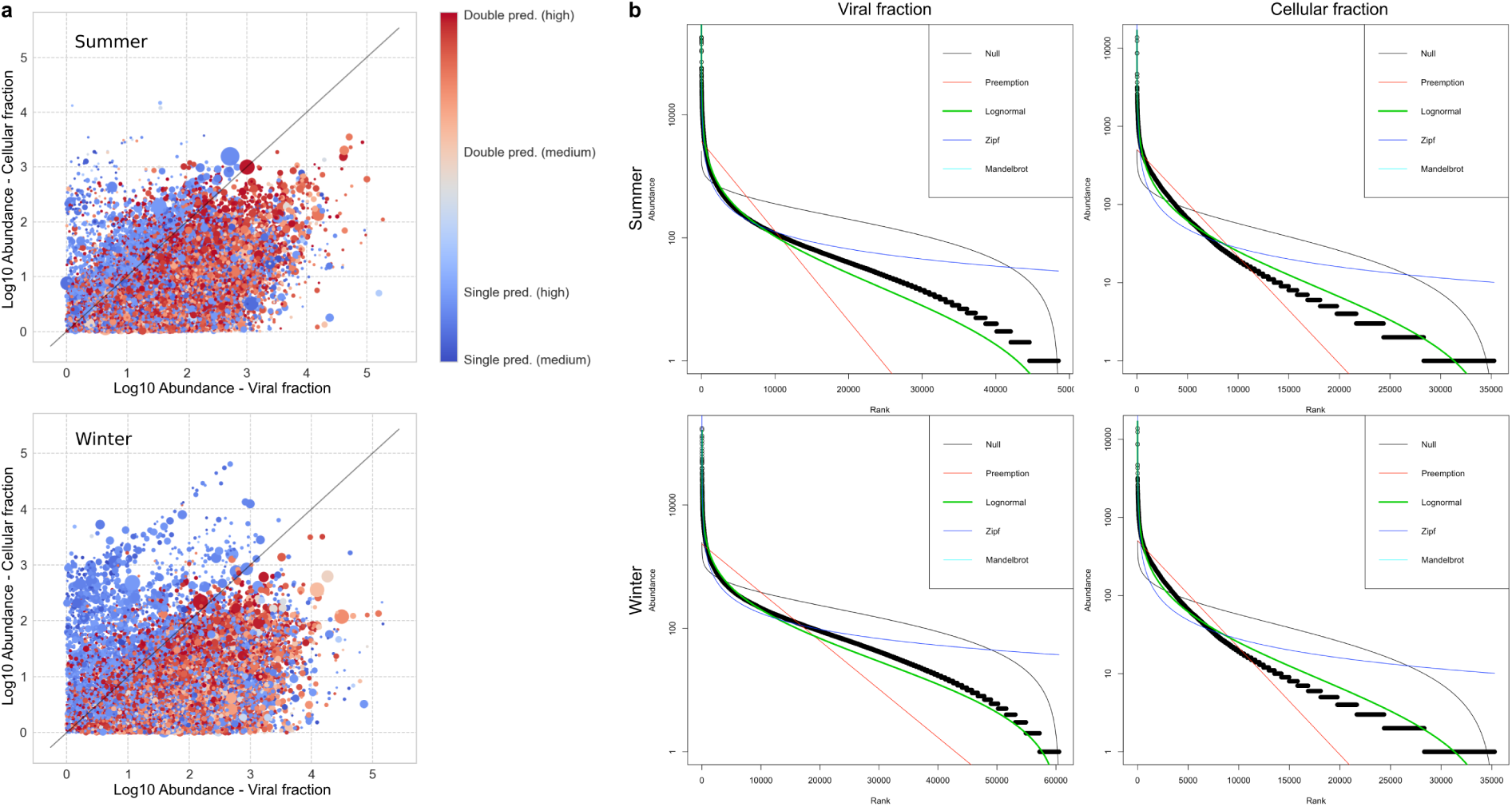
Viral contig prediction and abundance. (a) Abundance of contigs in the viral fraction (x-axis) and cellular fraction (y-axis) in the summer (top) and winter (bottom) samples. Colors scale from lower prediction score (blue) to higher score (red) as predicted by one or both virus prediction software VirFinder (89) and MARVEL (90). Size of each dot is proportional to the relative length of the contig. The diagonal line represents 1:1 abundance ratio. (b) Rank abundance dominance plots (RAD) of viral contigs (length ≥ 5kb and RPKM ≥ 1, open circles) by sample type and season. Fitted lines are different RAD models generated with vegan r package (34)

**Fig. S3.**
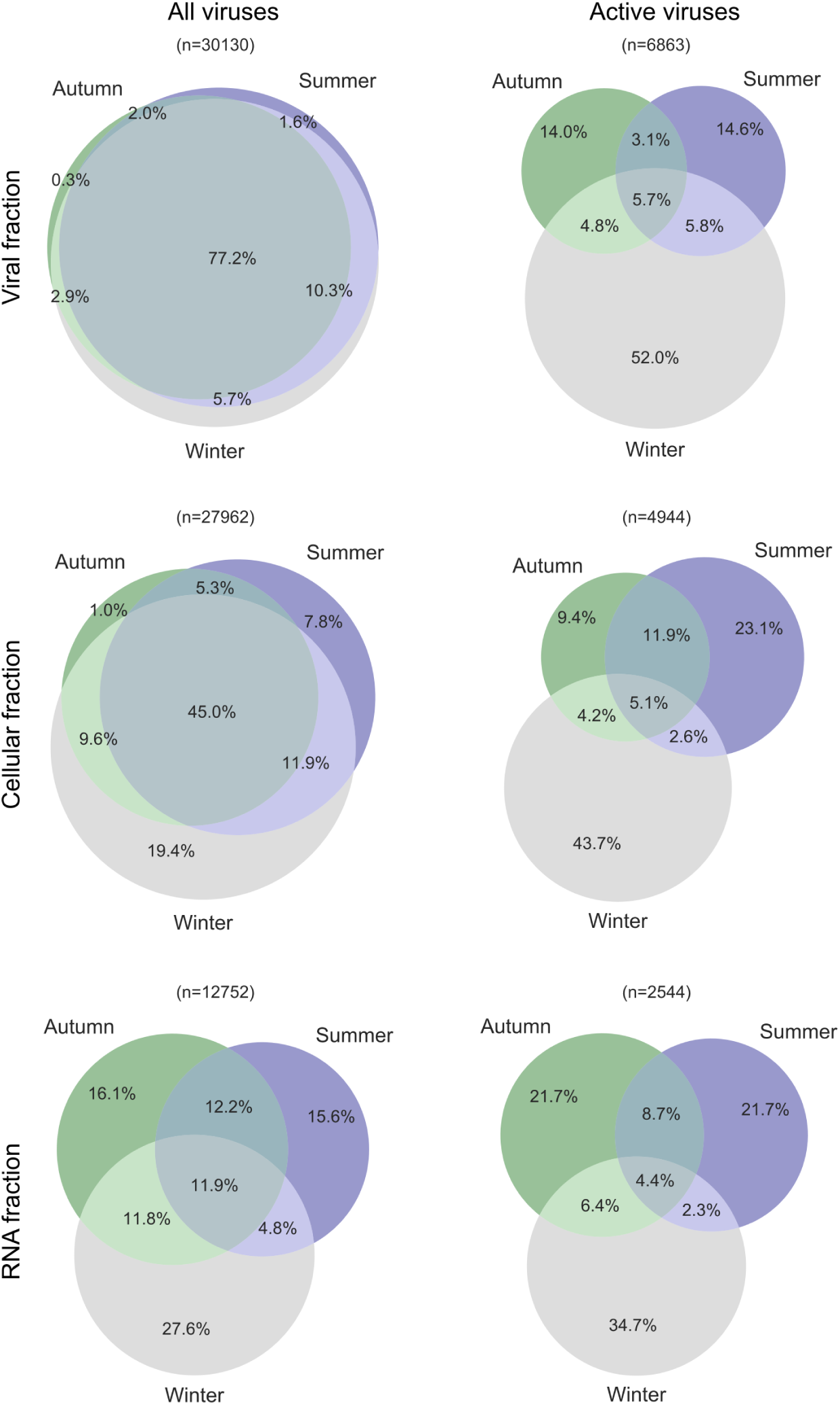
Percentage of viral contigs shared across seasons by sample type. Left: All viral contigs with count ≥ 1 RPKM in the virome, metagenome or metatranscriptome samples. Right: Most abundant viral contigs (Active group).

**Fig. S4.**
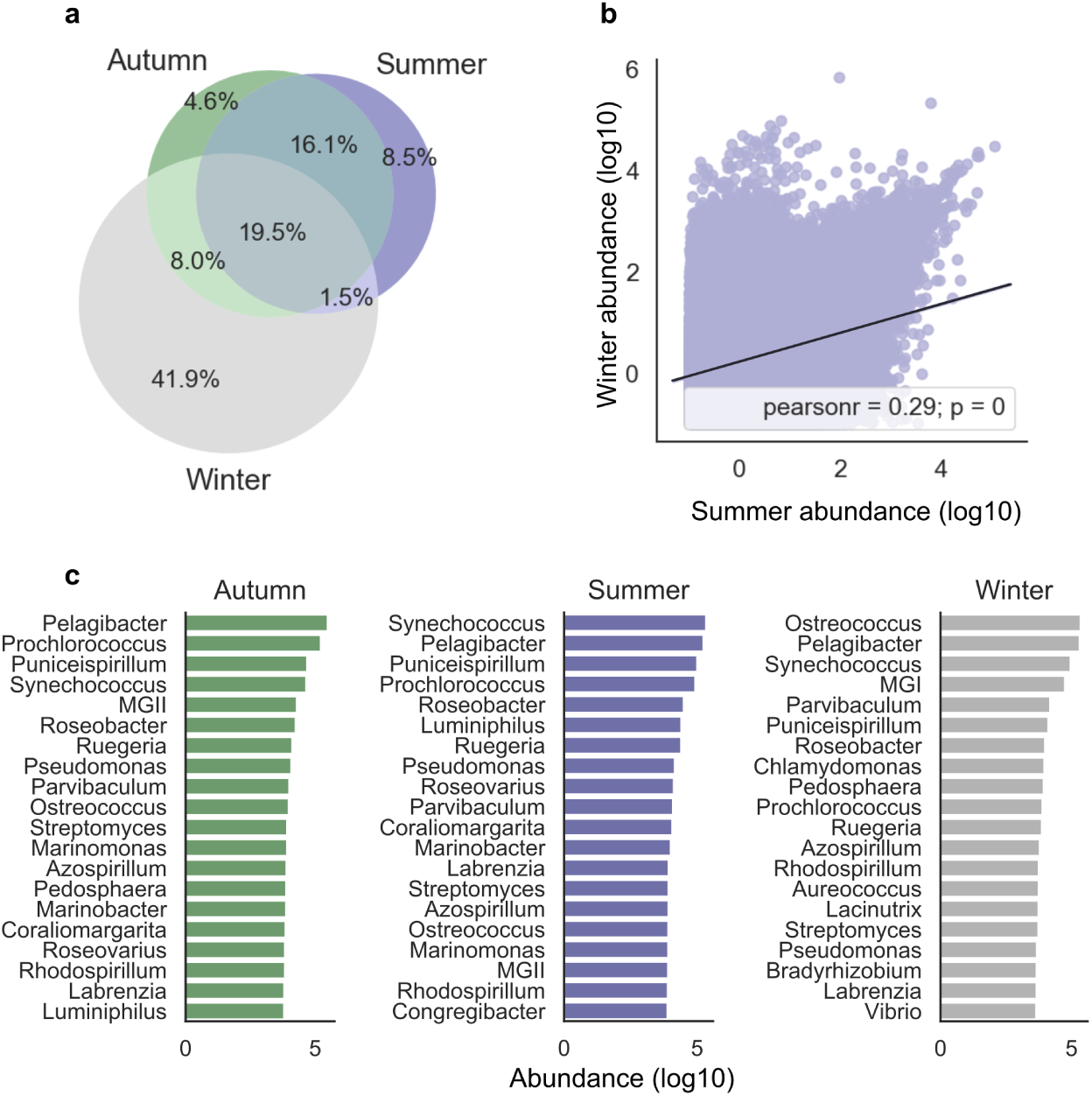
Microbial community seasonal abundance. (a) Percentage of microbial contigs shared between seasons with count ≥ 1 RPKM in the metagenome samples. (b) Abundance of microbial contigs in the summer vs. winter cellular samples. The fitted linear correlation line is shown for reference. (c) Most abundant microbial genera by season: autumn (green), summer (blue) and winter (grey).

**Fig. S5.**
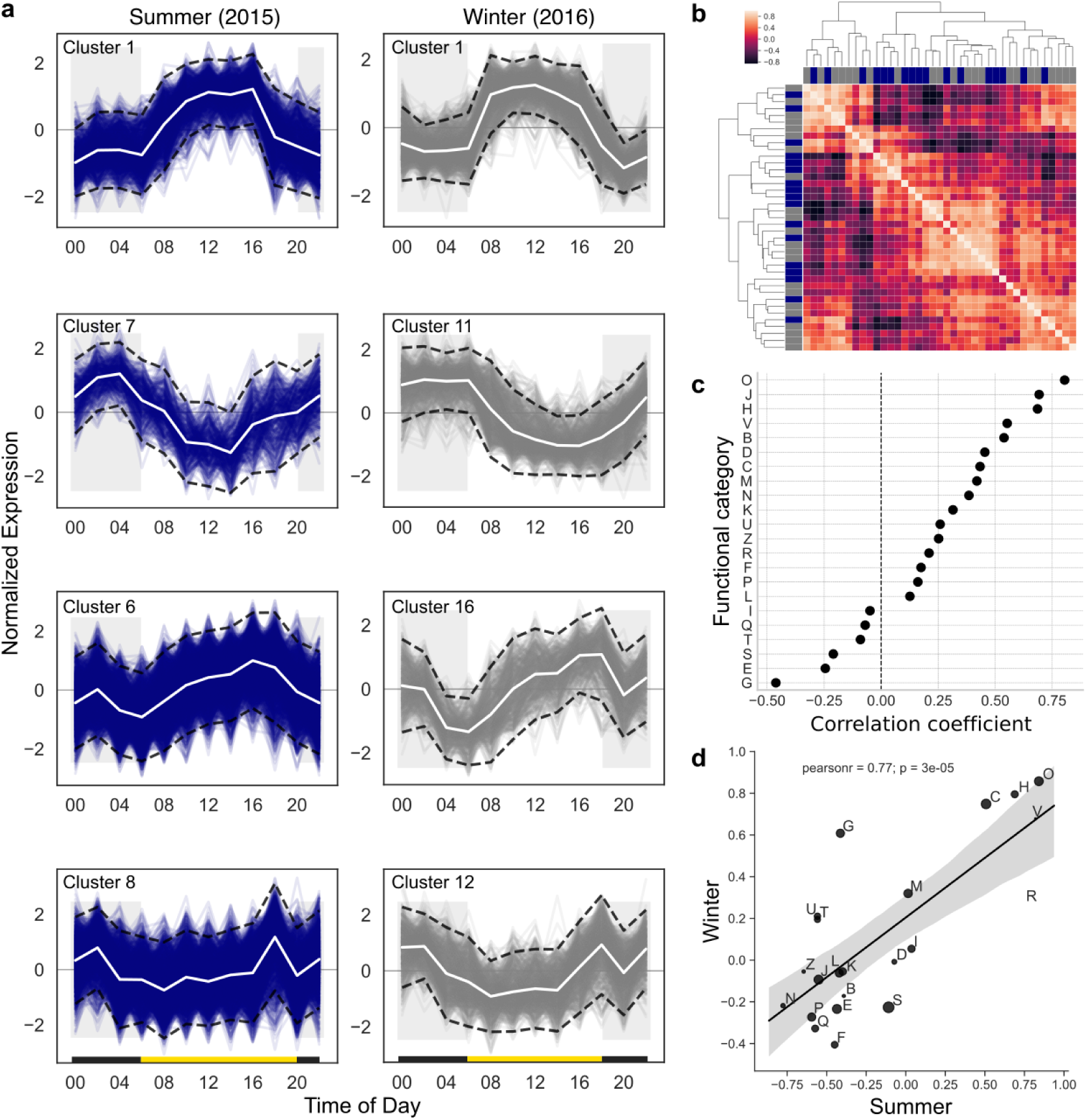
Microbial community expression patterns. (a) Selected gene clusters of similar expression patterns in the summer (blue) and winter (grey) metatranscriptomic samples. The genes were clustered using Gaussian mixture models coupled with Dirichlet process (DP-GP) (85). x-axis: time of day, y-axis: standard deviations from the mean, confidence bands (dashed lines) represent two standard deviations from the mean, horizontal bars: light (yellow) and dark (black) hours. (b) Heat map with hierarchical clustering of a pairwise correlation of the mean expression pattern of each of the DP-GP clusters. The similarity between the patterns is calculated based on Manhattan distance. Heat map colors range from light, indicating high similarity, to dark, indicating higher distance. Dendrogram tips are colored according to the season, summer (blue) and winter (grey). (c) Correlation of the mean expression by functional category between seasons. x-axis: Spearman R correlation coefficient, y-axis: COG functional categories (d) Correlation of functional categories with diel light intensity. x-axis: Correlation coefficient of summer functional groups, y-axis: Correlation coefficient of winter functional groups. The fitted linear correlation line is shown for reference.

**Fig. S6.**
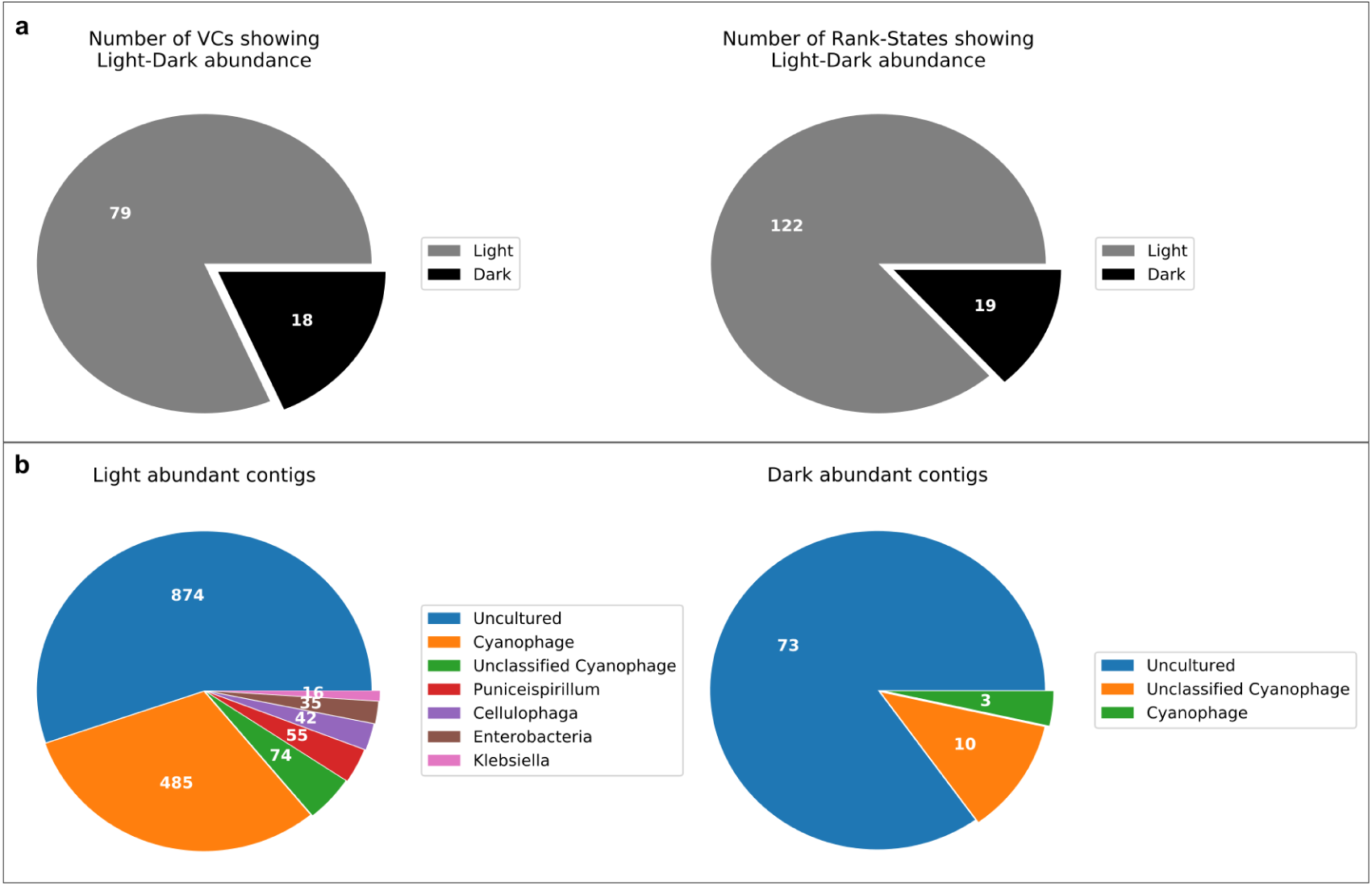
Light-dependent virus-host interaction (a) Number of viral clusters (minimum of 10 ORFs in cluster and minimum counts > 0 RPKM by ORF in either the light or dark samples), with at least one rank-state that displayed differential light-dark abundance (Mann-Whitney test < 0.05). (b) Number of contigs from 16 viral clusters and 9 rank-states (grouped by predicted host genus) with significant light-dark differential abundance (Mann-Whitney test < 0.05).

**Fig. S7.**
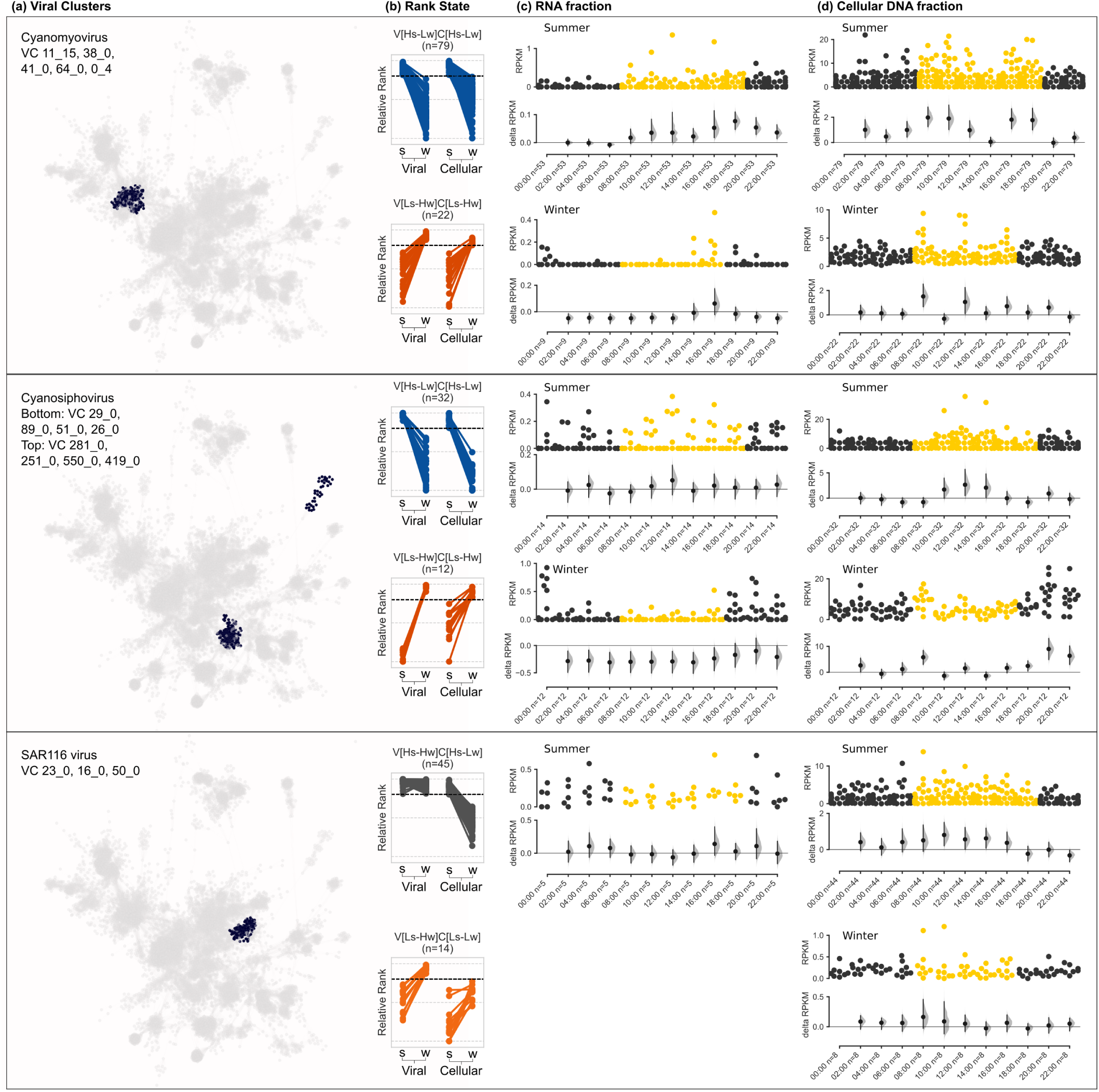
Intracellular diel abundance patterns of cyanomyoviruses, cyanosiphoviruses and SAR116 virus populations. (a) Viral clusters of contigs from three cyanophage families grouped using gene-sharing network (Bin Jang et al. 2019). (b) Seasonal rank-state abundance patterns of selected contigs from the VC on the left that displayed a differential light-dark signal (Mann-Whitney U test, p < 0.05). (c-d) Diel distribution of intracellular RNA (c) and DNA (c) of viral contigs from a specific VC and rank-state (left); x-axis, hours of the day when a sample was collected and the number of viral contigs expressed; upper panel, abundance distribution of viral contigs by time point (each point represents a contig); point color represent samples collected in the dark (black) and light (yellow); y-axis, contig abundance (RPKM); lower panel, estimation plot (37) displaying the effect size as a 95% confidence interval (1,000 bootstraps) of the mean differences between each time point compared against midnight as a reference group.

**Fig. S8.**
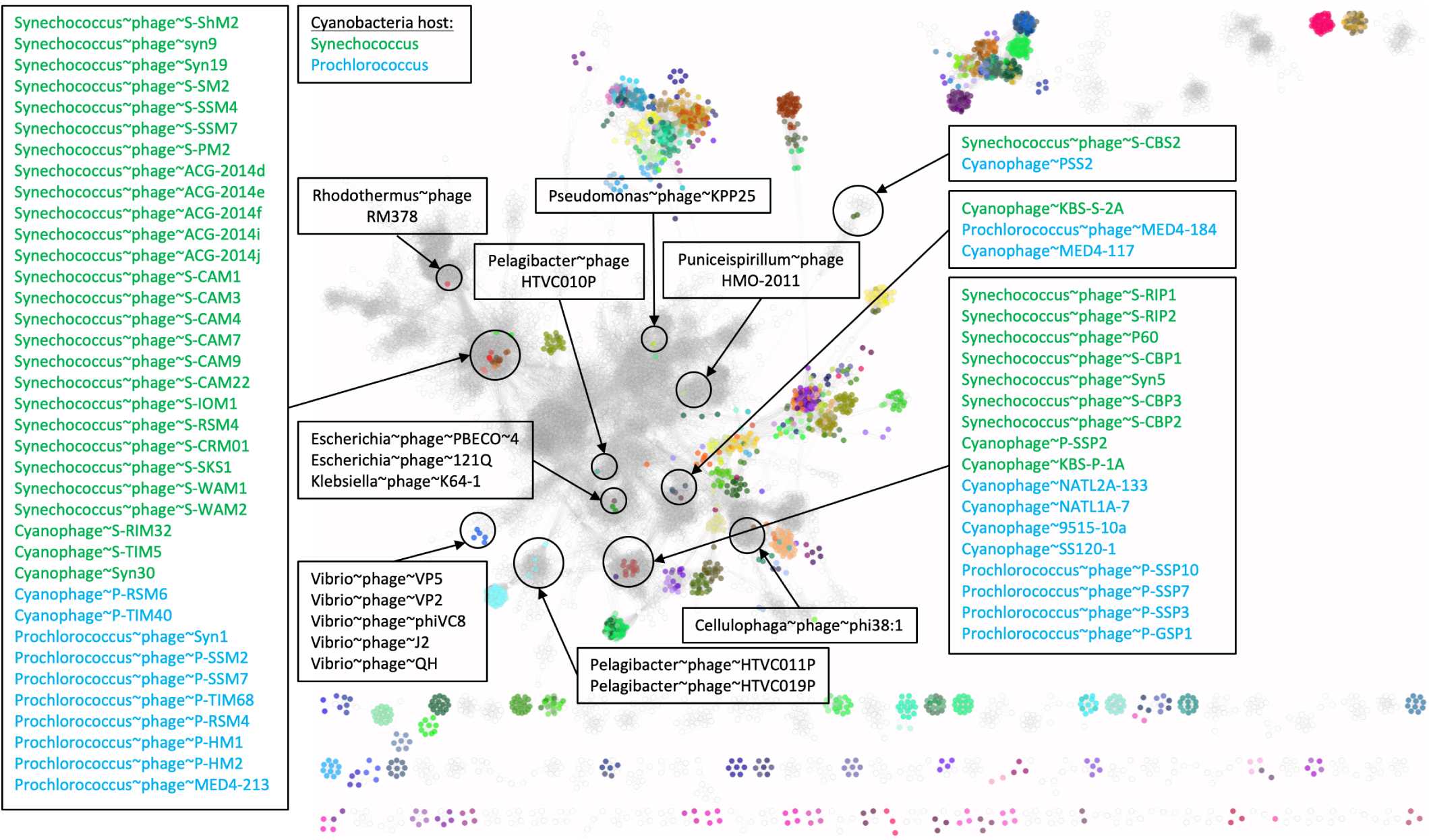
Gene sharing network of Refseq viral genomes and Red Sea predicted viral contigs (2,239 and 7,047 genomes, respectively). Grey nodes represent Red Sea viral contigs. Colored nodes represent RefSeq viral genomes. Labeled clusters represent taxonomic annotation based on Refseq viral genomes that appear in a specific cluster.

